# Coverage-dependent bias creates the appearance of binary splicing in single cells

**DOI:** 10.1101/2019.12.19.883256

**Authors:** Carlos F. Buen Abad Najar, Nir Yosef, Liana F. Lareau

## Abstract

Single cell RNA sequencing provides powerful insight into the factors that determine each cell’s unique identity, including variation in transcription and RNA splicing among diverse cell types. Previous studies led to the surprising observation that alternative splicing outcomes among single cells are highly variable and follow a bimodal pattern: a given cell consistently produces either one or the other isoform for a particular splicing choice, with few cells producing both isoforms. Here we show that this pattern arises almost entirely from technical limitations. We analyzed single cell alternative splicing in human and mouse single cell RNA-seq datasets, and modeled them with a probablistic simulator. Our simulations show that low gene expression and low capture efficiency distort the observed distribution of isoforms in single cells. This gives the appearance of a binary isoform distribution, even when the underlying reality is consistent with more than one isoform per cell. We show that accounting for the true amount of information recovered can produce biologically meaningful measurements of splicing in single cells.

## Introduction

Single-cell RNA sequencing (scRNA-seq) has provided impressive temporal resolution to our understanding of continuous biological processes such as cell differentiation [1, 2]. It has uncovered hidden heterogeneity among cells and exposed the factors that determine each cell’s unique identity. One broad source of transcriptomic diversity is alternative splicing, and several studies have uncovered compelling evidence of changes in alternative splicing among single cells during differentiation [3–6].

A particularly surprising conclusion of several scRNA-seq studies was the observation that splicing was often bimodal among supposedly homogeneous cells [5,7–9]. That is, some cells always spliced in a particular cassette exon, and some cells never spliced in the exon, but few cells showed truly intermediate inclusion within one cell. This unexpected result contrasted with previous single molecule imaging studies of several alternative exons that showed that cell-to-cell variability is minimized and tightly regulated by the splicing machinery in single cells [10]. This led to investigations of the mechanisms that might be responsible for stochastic splicing variability among apparently homogeneous cells, such as variation in DNA methylation [11].

We propose that the observed bimodality does not generally reflect splicing biology, but rather, that it exposes a technical limitation of the scRNA-seq data that have been collected so far. Because alternative isoforms of a gene share much of the same sequence, only the few RNA-seq reads mapping to the exact alternative splice junctions, or to the alternative exon itself, reveal its alternative splicing. When combined with the low mRNA capture efficiency of scRNA-seq and the PCR amplification of small amounts of starting material into a full-length sequencing library, these circumstances create the risk of bottlenecks that lose all but a few individual mRNAs of most genes in each cell.

The limitations of scRNA-seq are a known obstacle to studying splicing in single cells [12]. Similar concerns have arisen with the use of scRNA-seq to infer allelic expression; a careful analysis showed that stochastic patterns resulted from technical noise [13]. A recent study advised that the high dropout rate of scRNAseq makes it fundamentally unsuitable for measuring changes in splicing [14]. Others have implemented workarounds, e.g., using sequence features to predict splicing outcomes in lieu of sufficient sequencing coverage [6], or attempting to identify excess variance beyond technical noise [3, 11]. These studies have identified true examples of differential splicing in single cells, but they fundamentally do not explain how scRNA-seq limitations have caused qualitative, not just quantitative, distortions in our understanding of alternative splicing.

Here, we show that scRNA-seq splicing data are consistent with a simple model. Consider a particular cassette exon whose true pattern of exclusion follows a unimodal distribution of isoform ratios across cells (i.e., most cells express both isoforms, with a ratio revolving around the same mean). This distribution can be distorted by extreme information loss during library preparation and sequencing, creating the illusion that individual cells only produce one isoform or the other. Our simulations make it clear that the reliability of splicing measurements is a function of the initial amount of mRNA, the efficiency of its recovery, the underlying splicing rate, and further distortions from PCR amplification of cDNA. These effects should be considered when interpreting previous studies that used qualitative changes in the observed distribution of the splicing rates [5] or changes in their variance [11] as evidence for regulation of alternative splicing. Considering the true amount of information available for a cassette exon can allow for accurate observations of alternative splicing. Using a data normalization and filtering method to identify cassette exons with sufficient information, we are able to draw biologically relevant conclusions about alternative splicing in single cells.

## Results

We began by examining the splicing of several cassette exons in a high-coverage mouse scRNA-seq dataset [15], estimating their percent spliced-in as the fraction of splice junction reads that show exon inclusion (out of all reads that cover the junction). We use 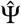 to denote these estimated rates, while Ψ denotes the actual rate as it is in the cell. For clarity we define 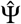 (which pertains to a specific cassette exon) as *binary* if it is close to 0 or 1 (i.e., the respective cell tends to express transcripts that either include the exon or exclude it, but not both). We further define the distribution of 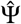 across cells as *bimodal* if its individual values are binary, where some cells have a 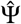 close to 1 (most observed transcripts include the exon) and some with 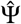 close to 0 (most observed transcripts do not include the exon). Strikingly, the exons we inspected had more binary outcomes in cells with fewer reads covering their splice junctions, while cells with more reads were more likely to show non-binary 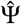 values (Figure 1a).This effect of coverage may reflect a nonbinary reality, since even if both isoforms are expressed in a certain cell, the likelihood of observing both isoforms is reduced as the number of captured mRNAs decreases. In contrast, if the underlying distribution was indeed bimodal, as previously proposed [5, 7, 8], then the read coverage would have little effect on the variability of 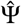 across cells.

**Figure 1:**
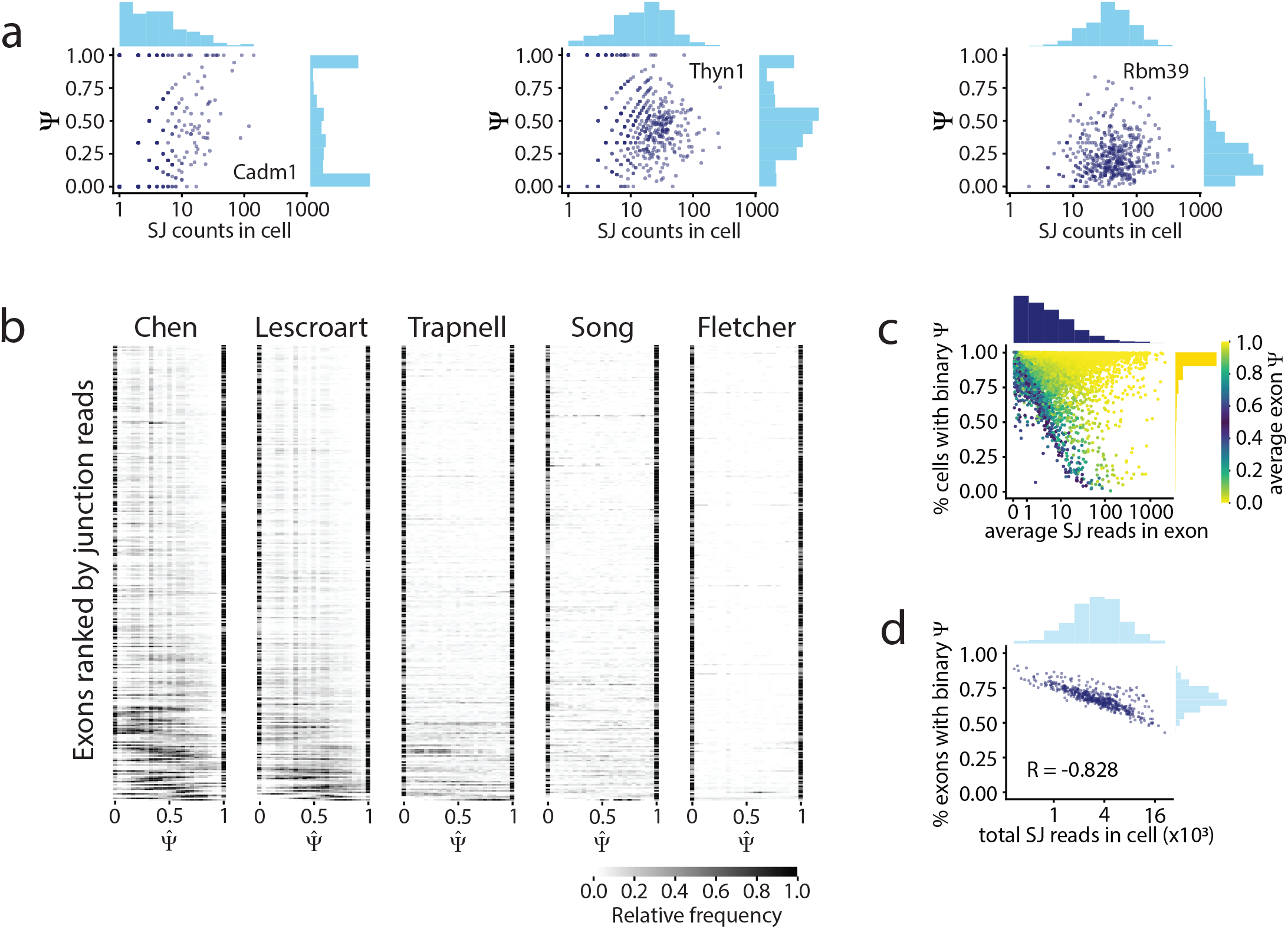
a) Comparison of splice junction read coverage and observed 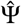 for three cassette exons in the Chen dataset, with low, medium, and high coverage [15]. Each dot represents the 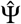 of that exon in one cell. b) 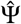 distribution of the 300 highest coverage cassette exons with intermediate splicing (average 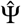 between 0.2 and 0.8) in each of the five analysed datasets (Table 1). Each row in the heatmap shows the distribution of 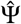 for one exon across all cells. c) Relationship between the average read coverage and proportion of binary observations for each cassette exon in the Chen dataset. d) Correlation between the total number of splice junction reads captured in each cell, and proportion of cassette exons that show binary 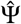 in that cell.

To further explore this phenomenon, we extended our analysis to the entire observed transcriptome in several full-transcript scRNA-seq datasets from mice and humans. We found a strong effect of coverage on the observed bimodality in all cases. We consistently found that the bimodally distributed 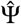 values were mostly observed in junctions with low coverage (Figure 1b,c; Supplementary Figure S1a,b). We find similar results in an alternative analysis whereby cells with an overall higher number of splice junction reads also tended to have a smaller fraction of exons with binary values (Figure 1d, Supplementary Figure S2c,d). Interestingly, in a closer inspection, we find that the association between binary values and read coverage is not observed in exons that are binary but not bimodal (i.e., the average 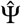 is close to 0 or 1; Figure 1c). Taken together, these observations suggest that the presence of bimodality in exon inclusion patterns may reflect a distortion of an underlying unimodal splicing distribution (i.e., when cells in fact express both isoforms), rather than a truly bimodal expression pattern in the analyzed cells.

A simple probabilistic exercise shows the potential loss of splicing information during sequencing. Single cell RNA-seq experiments that capture full-length transcripts have an estimated capture efficiency of only 10%, due to RNA degradation and inefficient reverse transcription [4, 8, 16, 17]. For instance, a gene that expresses 20 mRNA molecules in a cell might only have two mRNAs recovered, and if that gene is alternatively spliced with a true splicing rate Ψ of 0.5, there is approximately a 50% chance that those few recovered mRNAs will only represent one of the two isoforms that were originally present in the cell (Supplementary Figure S2a,b). As many genes are expressed at just a few RNA molecules per cell, low recovery might affect many alternative splicing events [18, 19]. Furthermore, while the empirically observed 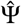 provides a maximum likelihood estimate for the true splicing rate, the uncertainty of this estimate (i.e., the range of alternative values with a nearly similar likelihood) decreases substantially with the number of observed molecules (Supplementary Figure S2c and Methods).

### Simulations of RNA sequencing reveal technical sources of distortion of splicing estimates

Our theoretical reasoning above relied on a simple model where the number of observed mRNA molecules (rather than number of reads) is known and the only distorting factor is a limited capture efficiency. In practice, both of these assumptions are challenged due to additional factors, such as PCR amplification and variability in the capture efficiency across cells. To investigate the pertaining effects on 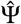 distributions in this more complex setting, we designed a probabilistic simulator of alternative splicing in single cells (Figure 2a). The model has two main components: We begin by simulating the underlying molecular content of each cell, by drawing gene expression levels and cassette exon splicing rates from a probabilistic model of cell state. We then simulate the technical process of extracting data from each cell using single cell RNA sequencing with a full transcript coverage. This part accounts for variability in capture rates, and the effects of PCR amplification, fragmentation and sequencing. It relies on SymSim, a simulation software for single cells RNA sequencing data [20]. The final product of our simulation is the number of splice junction reads that either span or skip each exon in each cell. These numbers are distorted in a way that reflects real nuisance factors. For instance, two reads could have originated from the same molecule due to amplification effects.

**Figure 2:**
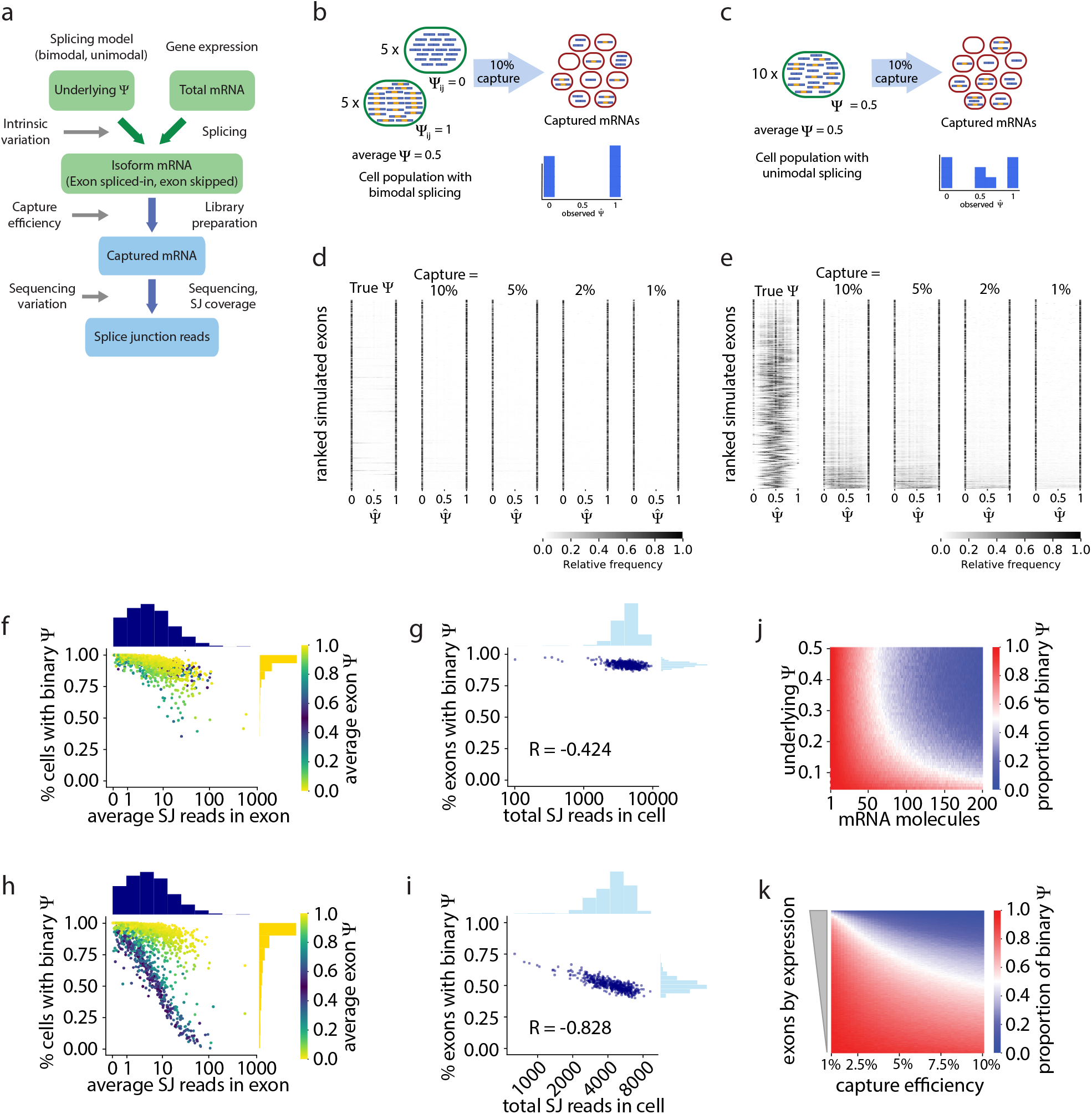
a) Simulator of scRNA-seq splicing data. Green elements represent biological variables; blue elements represent technical processes. The underlying Ψ is drawn from a Beta distribution with either a bimodal or unimodal shape. Individual gene expression determines the total number of mRNAs, and these mRNAs are spliced stochastically according to Ψ, producing isoforms that splice in or skip the exon. mRNAs are captured with a probability drawn from a normal distribution. Sequencing produces splice junction reads, which determine the observed 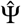. b) Bimodal model of cassette exon splicing. Some cells consistently splice in the exon, while others consistently skip it. After mRNA capture and sequencing, observations of Ψ are almost exclusively binary. c) Unimodal model. Individual cells express some mRNAs that splice in the cassette exon and some that skip it. Low mRNA capture dramatically reduces the number of cells in which both isoforms are observed, artificially inflating binary Ψ values. d) Simulations of alternative splicing and scRNA-seq under the bimodal model. The observed 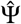 distribution is similar to the true Ψ distribution, and its shape is largely unaffected by capture efficiency. e) Simulations with the unimodal model. Exons with high expression have a unimodal distribution of true Ψ. The observed distribution of 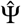 is distorted by low capture efficiency, and only a handful of the highest expressed exons maintain a unimodal 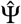. Fewer exons show unimodal splicing as the capture efficiency is reduced. f) Under the bimodal model, exons with high coverage have slightly fewer binary Ψ observations, and g) simulated cells with a high number of total splice junction reads have slightly fewer exons with binary 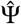. h) Under the unimodal model, exons with intermediate splicing show a strong decrease in binary observations as coverage increases, as seen in real data (Figure 1c). i) Similarly, simulated cells with high read coverage have a decrease of the proportion of binary 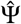. j) Effect of the initial number of mRNA molecules and underlying Ψ on the proportion of binary 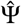 observations. k) Effect of capture efficiency on the proportion of binary observations of cassette exons with underlying Ψ = 0.5.

We used our simulator to investigate how the observed inclusion (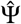) of cassette exons differs from the underlying Ψ, setting the average capture rate to 10% and the other technical parameters to values that are characteristic of Smart-seq2 datasets (see methods). We considered either a bimodal and binary regime of Ψ (i.e., both isoforms are expressed in the population, but rarely by the same cell; Figure 2b), or a nonbinary regime (cells tend to express both isoforms; Figure 2c) [5, 11, 21]. We simulated the splicing of cassette exons in 500 genes, in a population of 300 single cells.

As expected, in the bimodal case, the observed estimates (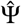) reflected the underlying process well, independent of the average capture efficiency (Figure 2d). In contrast, when we modeled a non-binary splicing regime, the observed 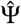 distributions were strikingly similar to the splicing distributions of cassette exons in real single cell RNA-seq datasets (Figure 1b, 2e). Specifically, the loss of information due to mRNA recovery and library generation led many of the observed 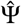 to become binary, and their distribution across cells to become bimodal. This tendency again correlated with coverage, whereby lowly covered exons showed the strongest effect, while exons with high coverage maintained a non-binary distribution. Consistently, in this non-binary regime, the average of 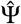 was similar to the true average of Ψ, but the variance of 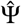 increased (Supplementary Figure S2d,e). Furthermore, as in the real data sets (Figure 1c), we also found that the dependency between read coverage and the chance of observing a binary 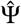 is more pronounced in exons with an underlying Ψ that is far from binary (Figure 2f-i), highlighting again that such an association likely indicates an artifact.

Finally, we estimated the chance of observing only one type of isoform (i.e., a binary 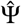) as a function of the underlying Ψ and the number of transcripts that are present in the cell. As above, our simulations assume an average mRNA capture rate of 10% Figure 2j). Our results delineate the range of values in which an artifact is less expected. For instance, for an exon with 50% inclusion rate, a non binary estimate is more likely if the respective gene has at least 50 transcripts in the cell. Notably, these estimates are more conservative than the theoretical analysis (Supplementary Figure S2a,b), due to the effect of the technical nuisance factors we modeled in this simulation.

#### Technical sources of variation in scRNA-seq

To address the extent of the distortion from technical factors, we ran the simulator with a fixed underlying Ψ = 0.5, and observed the effect of varying different parameters of the RNA sequencing process. Decreasing the average capture efficiency dramatically increased the number of binary 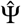 observations, particularly for exons with low expression, although even highly expressed genes suffered great distortion in the observed distribution of 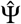 when the average capture efficiency was very low (Figure 2k). These results reinforced the hypothesis that capture efficiency is a main technical factor that creates the appearance of bimodality in single cell splicing.

Another potential source of technical variation is the amplification process. To generate the sequencing libraries in studies of splicing in single cells, mRNAs are usually reverse transcribed, amplified as full-length cDNAs, fragmented, and finally amplified again to generate enough material for sequencing. Previous studies have proposed increasing the number of PCR cycles for cDNA amplification to compensate for the low starting material in scRNA-seq [5, 14]. Here, we find that increasing the number of cDNA amplification PCR cycles resulted in a modest, yet consistent increase in the number of binary 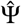 observations (Supplementary Figure S2f,g), which might explain the differences between the theoretical and the simulation-based analysis. Notably, the PCR effect was modestly alleviated by increasing the likelihood of duplicating a cDNA molecule every PCR cycle (i.e., amplification efficiency [20]; see Supplementary Figure S2f).

### Exon filtering enables a more accurate analysis of splicing in single cells

We next set out to find criteria that would identify cassette exons with enough information to draw biological conclusions about their splicing. While our filter should ideally rely on the actual number of mRNA molecules observed, full-length RNA-seq experiments generally do not report an absolute mRNA count. Previous studies used filters based on the number of reads covering alternative splice junctions as a proxy for the amount of information [5, 11]. However, the number of splice junction reads is influenced not only by the number of recovered mRNAs, but also by the extent of PCR amplification and sequencing depth.

To estimate the number of mRNA molecules that were captured into cDNA, we adapted the Census normalization approach [4]. This method infers a per-cell scaling factor between the relative abundance of each gene, inferred from RNA-seq, and the actual number of mRNAs recovered (Figure 3b). We found that some datasets with many reads per cell, such as the Song et al. dataset [5] (Supplementary Figure S3b), nonetheless had few mRNAs recovered per cell, which may explain the extensive splicing bimodality in this dataset. The dataset with the highest recovery of mRNAs (Chen et al [15]) indeed showed a larger extent of non-binary splicing events.

**Figure 3:**
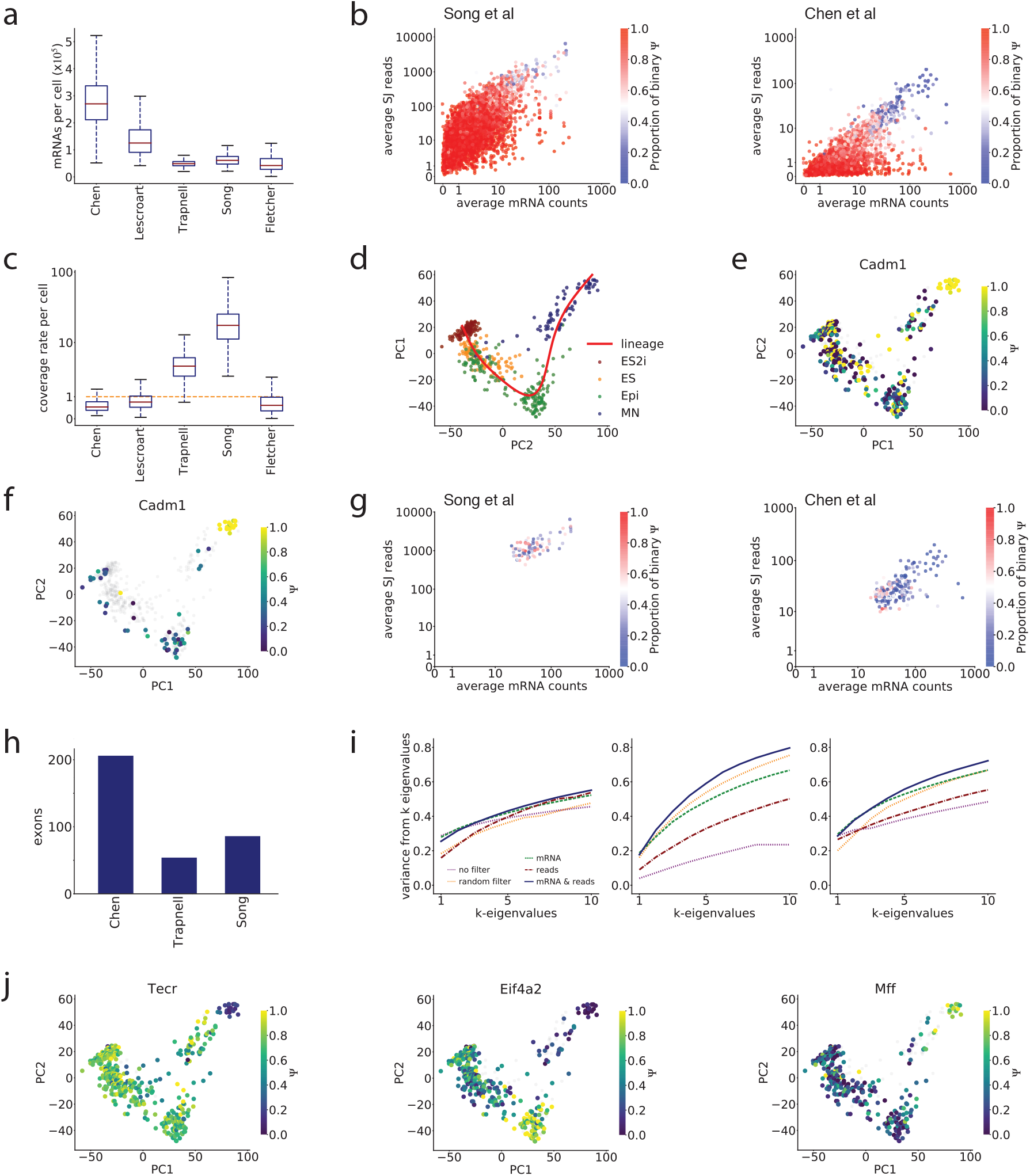
a) Total mRNA molecules captured per cell in each dataset, estimated with the Census normalization [4]. b) The estimated number of recovered mRNAs and the number of splice junction reads for each cassette exon, averaged across cells, for the Song et al and Chen et al datasets. c) Per-cell coverage rate in each dataset, showing how many reads are expected to cover each position of an mRNA. d) PCA projection and inferred lineage of single cells in the Chen dataset based on gene expression, showing differentiation of mouse ES cells into motor neurons. e) Cadm1 alternative splicing appears binary in many cells, with little distinguishable pattern. f) After removing cells with fewer than 10 recovered Cadm1 mRNA molecules and fewer splice junction reads than expected from 10 mRNAs (grey), a clear pattern of differential splicing during differentiation is visible. g) Cassette exons were filtered to remove observations in individual cells with fewer than 10 recovered mRNAs, or with fewer splice junction reads than expected from 10 mRNAs; we then kept exons that passed these filters in at least half of cells. The plot shows the average number of recovered mRNAs and average number of splice junction reads for only the remaining observations, for each exon that passed the overall filter. The remaining exons have fewer binary 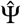 observations. h) Number of exons passing the filter for three datasets. i) Comparison of covariance of 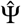s using three methods of filtering Ψ observations (exons with more than 10 mRNAs in a cell, exons with more than 10 splice junction reads in a cell, and the data-driven filter described in panel g), as well as two controls (no filter, and a random subsample of the data to match the number of observations that pass the mRNA + reads filter). Covariance structure was measured as the sum of the squares of the first *k* eigenvalues of the covariance matrix, divided by the sum of all squared eigenvalues. j) Example exons that pass the overall filtering criteria in the Chen dataset. Clear patterns of splicing change are observed in the cells that individually pass the filter (color); cells that do not meet the criteria are not considered (grey).

Next, we explored how many mRNAs must be recovered to reduce the prevalence of binary observations. Our simulations showed that a median of nine recovered mRNA molecules in a cell were required to give a 50% chance of observing both isoforms of an exon with an intermediate inclusion level (Figure 2j, Supplementary figure S3f). In keeping with this, we saw in the real data that exons with an average of at least 10 mRNAs recovered per cell generally had substantially fewer binary observations (Figure 3b, Supplementary Figure S3c-e). However, a subset of these exons had few splice junction reads relative to the estimated mRNA count, and this subset showed binary splicing in many cells. We expect that these represent cases where the overall read coverage of the full gene led to a high recovered mRNA estimate, but fewer reads were recovered from the specific splice junctions of interest, perhaps due to annotation errors or poor recovery of fragments with particular sequence composition.

To prevent distortions arising from this low read recovery, we considered the number of splice junction reads expected to arise from a cassette exon. First, for each cell we calculated the coverage rate, representing the expected reads per position of an mRNA, as the total nucleotides of reads divided by the total nucleotides of mRNAs recovered from that cell (Figure 3c). Then, the expected number of splice junction reads for a particular cassette exon also depends on its Ψ: with no amplification, one mRNA molecule will produce one pertinent splice junction read if the cassette exon was excluded, or two if it was spliced in. Thus, 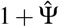 splice junction reads are expected per mRNA, multiplied by the per-cell coverage rate. This provided a second filtering criterion: we excluded exons with fewer reads than would arise from 10 mRNAs, given that exon’s 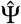 and the coverage rate in that particular cell. This metric is calculated for each exon separately, driven by the actual information in each cell.

Our filter based on the number of recovered mRNA molecules and the expected splice junction reads was able to remove potentially spurious binary observations, as shown for alternatively spliced exon 8 of Cadm1, a cell adhesion factor that is differentially spliced in neurogenesis [22, 23]. Single-cell data showed an apparent bimodal distribution of Cadm1 splicing throughout differentiation, with no clear pattern of differential splicing (Figure 3d,e). However, discarding cells that had fewer than 10 mRNA molecules and fewer splice junction reads than expected from 10 mRNA molecules revealed a clear change in Cadm1 splicing during neurogenesis (Figure 3f).

Finally, we asked if these two filtering criteria would allow us to identify exons with sufficient alternative splicing information across many cells. We screened all exons with average 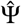 between 0.05 and 0.95, and selected the exons in an individual cell that passed both filters. We further required that the exon pass these cutoffs in at least half of cells. This simple approach selected exons that were predominantly unimodal (Figure 3g), and we hypothesized that the remaining exons with bimodal splicing represented legitimate, regulated changes in splicing between cells. These exons should show some co-regulation, which should be reflected in the covariance of the 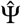s. We compared the structure of the covariance matrix of cells using the exons passing our filter with the covariance matrices of cells with exons passing two other selection criteria: a cutoff of 10 recovered mRNAs without considering the splice junction reads, and a simple read-based cutoff of 10 splice junction reads as used in many previous analyses (Figure 3h,i). Our combined filter recovered more evidence of co-regulation than the simple read-based filter. Importantly, many exons that passed our filter showed clear patterns of differential splicing between cell subtypes, including some with known regulation during differentiation (Figure 3j) [23]. We concluded that scRNA-seq can reveal true changes in alternative splicing, including truly bimodal distributions between cell types, when noisy observations are filtered out by principled coverage criteria.

## Discussion

The surprising result that alternatively splicing is bimodal among single cells provoked curiosity and speculation. Bimodal outcomes might reflect hidden cell subtypes, but the bimodality was seen even among apparently homogeneous cells. Did splicing outcomes reflect some unknown, stochastic cell state?

We have shown here that the bimodal patterns could have an entirely different explanation: profound technical limitations of single cell RNA sequencing. A crucial limit on biologically meaningful splicing observations in a single cell is the number of mRNAs available to inform the measurement. This is determined both by the expression of the genes that contain the exons of interest, and by the capture efficiency of the experiment. It is important to note that the depth of a sequencing library does not necessarily reflect its quality. Along with low mRNA numbers, splicing observations are also distorted by uneven amplification efficiency and cDNA overamplification. Increasing PCR amplification cycles in an attempt to compensate for low capture efficiency has the risk of worsening the technical distortion. Indeed, in our analysis, the dataset with the highest read count per cell actually had quite low mRNA recovery and large technical distortion, creating an appearance of bimodal splicing [5]. Moreover, a qualitative change in the observed 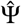 distribution of an exon between single cell subpopulations does not necessarily reflect a change in the underlying splicing rate, as changes in gene expression and mRNA recovery between samples can create the illusion of a splicing change.

Further developments in statistical analysis that carefully account for both missing and redundant information due to low capture efficiency could make splicing observations in single cells more reliable. We set the foundation for such analysis by proposing a probabilistic process that describes the biological and technical steps that generate single cell splicing data. We also introduced a simple approach that builds on the Census normalization [4] to estimate the number of mRNAs recovered and the extent of artificial duplication of splicing information. This metric provides a practical filter for identifying exons with sufficient information to analyze. On the experimental side, improving the capture efficiency of scRNA-seq methods while moderating the extent of overamplification is crucial for increasing the subset of exons for which reliable observation can be made.

True biological insight into alternative splicing can indeed be found from high-quality scRNA-seq data, and we hope that new methods will allow better understanding of splicing regulation, cell-to-cell variation, and the importance of alternative splicing in defining cell fate [24, 25]. However, some limitations are inherent to the situation. Single cells express a limited number of mRNAs per gene; splicing observations in single cells will always be inherently noisy reflections of the underlying biology.

## Methods

Analysis code is available at https://github.com/lareaulab/sc_binary_splicing

### Analysis of single cell RNA-seq datasets

#### Datasets

Five publicly available single cell RNA sequencing datasets were analysed (Table 1). These datasets are referenced with the first author’s lastname throughout this paper. For the Chen dataset, we only selected the cells that are part of an experiment that induced mouse embryonic stem cell differentiation to motor neurons (total 488 out of 617). For the Trapnell dataset, we only selected the runs that are annotated to have one cell per well, as opposed to zero or two (314 out of 372 single cell samples).

**Table 1:** Single cell RNA-seq datasets.

| Dataset | Organism | Biological process | no. cells | No. ASE | reads/event | Accession | Ref. |
| --- | --- | --- | --- | --- | --- | --- | --- |
| Chen | mouse | mES motor neuron differentiation | 488 | 1726 | 38.4 | GSE74155 | [15] |
| Lescroart | mouse | cardiomyogenesis | 598 | 1808 | 29.5 | GSE100471 | [36] |
| Trapnell | human | skeletal myogenesis | 314 | 1053 | 98.0 | GSE52529 | [33] |
| Song | human | iPS motor neuron differentiation | 206 | 971 | 703.0 | GSE85908 | [5] |
| Fletcher | mouse | olfactory neurogenesis | 849 | 287 | 8.8 | GSE95601 | [37] |

#### Alignment, TPM quantification and Ψ estimation

We aligned the reads of each dataset using STAR 2.5.3 [26] with two-pass mode and index overhang length adapted to the read length of each dataset. We used the hg38 genome annotation for the human RNA-seq datasets, and the mm10 annotation for the mouse datasets. Gene expression levels in transcripts per million (TPM) were calculated by running RSEM [27] on the BAM files produced by the STAR alignment. We ran rMATS 3.2.5 [28] on bulk human and mouse RNA-seq datasets from cell types matching the scRNA-seq datasets [15, 29, 30] to find all annotated cassette exon alternative splicing events in each cell type. Then we used the SJ.out.tab files obtained from the scRNA-seq STAR alignment to obtain the splice junction reads compatible with the list of cassette exons found by rMATS. For each cell *i*, we calculated the observed Ψ of the cassette exon *j* as:

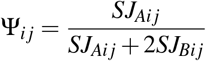

where *SJ*_*Aij*_ correspond to the number of reads that cover the two splice junctions compatible with cassette exon inclusion, and *SJ*_*Bij*_ are the reads that cover the splice junction compatible with its exclusion.

We also determined the coverage of an exon *j* in *i* as *SJ*_*ij*_ = *SJ*_*Aij*_ + *SJ*_*Bij*_. We used *SJ*_*ij*_ and Ψ_*ij*_ for the analyses shown in Figure 1.

#### Gene expression normalization and pseudotime inference

For the purpose of visualization of the Chen et al. [15] dataset as shown in Figure 3, we normalized the gene expression data. First we selected the genes with TPM> 5 and more than 20 reads in at least 20 cells. After filtering, we used SCONE 1.6.1 [31] to select the best normalization approach for the data. For improving the normalization of the data, we used additional information for each cell, including the annotated cell type and batch, total number of reads, housekeeping genes and genes that are expected to change in the biological process that the dataset covers. We applied principal component analysis (PCA) over the log-counts from the best SCONE normalization, and used the first two principal components to infer pseudotime using Slingshot 1.0.0 [32]. We used the cell type annotation as the cluster input for slingshot, and manually indicated the direction of the biological process.

#### mRNA counts estimation with the Census approach

We performed our own implementation of the mRNA count estimation proposed by Qiu et al in Census [4]. The total number of transcript mRNAs in cell *i* is estimated as:

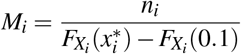

where 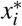 is the mode of the log-transformed distribution of TPM values in cell *i*. As in Qiu et al, we found 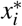 by fitting a Gaussian kernel density estimation to each distribution and finding its peak. We also set 0.1 as is the minimum TPM below which it is assumed that no mRNA is present. *n*_*i*_ is the number of genes in cell *i* with an estimated TPM in the interval 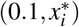. 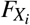 is the cumulative distribution function of the TPM values in cell *i*. The original Census implementation also adjusts the mRNA estimation by multiplying by 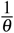, where *θ* is the capture efficiency of the dataset estimated with RNA spike-ins. Since for most datasets we do not have a reliable way of estimating the capture efficiency, we removed this adjustment from the equation, so that *M*_*i*_ in our estimation is not an estimation of the amount of mRNAs present in the cell lysate as it is in Census, but an estimate of the mRNAs successfully captured into cDNA.

We found that some datasets contained outlier cells with *M*_*i*_ much higher than the median estimate (more than ten-fold increases). These outliers generally correspond to cells with a multimodal TPM distribution. An inflation in very low TPM values distorts the normalization by shrinking the values of 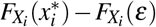, thus inflating the *M*_*i*_ values in these cells. Because the Census method relies on a Gaussian kernel density estimation that performs inaccurately for multimodal distributions, we excluded this handful of outliers from further analysis.

Finally, the number of mRNA transcripts of gene *g* in cell *i* is calculated as:

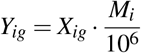

where *X*_*ig*_ is the expression of gene *g* in cell *i* expressed in TPM.

#### Nucleotide coverage and expected splice junction reads

Amplification in short-read library preparation can lead to multiple reads from different cDNA copies covering the same nucleotide from a single mRNA molecule. We estimated the coverage rate of each cell as the expected number of reads covering a single nucleotide as:

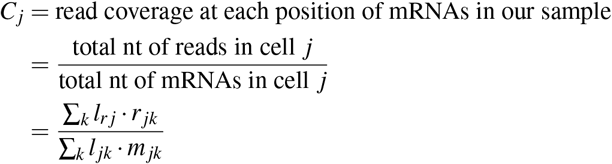

where *C*_*j*_ is the number of reads expected to cover each nucleotide in cell *j*; *l*_*r*_ is the effective read length for the sequencing protocol used in cell *j*; *r*_*ij*_ is the number of reads that map to gene *i* in cell *j*, as reported by RSEM; *l*_*ij*_ is the effective length of gene *i* in cell *j*, as reported by RSEM; and *m*_*ij*_ is the estimated number of captured mRNAs of gene *i* in cell *j*.

We assume that in each cell, the expected number of reads covering a splice junction is the same as the number of reads expected to cover each nucleotide. We also assume that this expected number is uniform across all mRNAs in the cell. We do not consider other factors that might affect this coverage such as sequencing biases. Therefore, the expected number of splice junction reads covering the splicing of cassette exon *i* in cell *j* is estimated as:

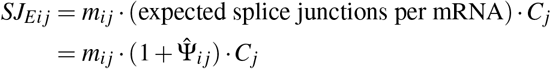

where *SJ*_*Eij*_ is the expected number of splice junction reads covering the splicing of exon *i* in cell *j* (both for mRNAs that splice-in or skip the exon). *m*_*ij*_ is the estimated number of mRNAs from the gene containing the cassette exon *i* in cell *j*; 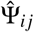 is the observed splicing rate of exon *i* in cell *j*. The expected number of splice junctions per mRNA is 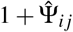 because 1 splice junction read is present in mRNA molecules that skip the exon, and 2 in those that include it.

#### Filtering and covariance structure analysis

Simulations of the effect of the initial number of mRNA molecules of a gene and the underlying Ψ suggest that, an average of 61.81 mRNA molecules are necessary to have a 50% chance of making an intermediate Ψ observation when the underlying Ψ is 0.5. This number goes up to 89.78 if the Ψ is 0.2 or 0.8 (Supplementary Figure 3f). Assuming a capture efficiency of 10%, we rounded at 10 captured mRNA molecules as the lower threshold for a quality Ψ observation.

In some cases, the number of observed splice junction reads is discordant with the estimated number of mRNAs recovered. Therefore we set a additional filter based on the number of reads expected to come from 10 mRNA molecules that are informative about the splicing of a cassette exon:

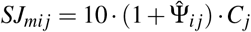

Therefore, for every observation, we required at least 10 mRNAs of the gene captured, and at least the number of reads that we expect if 10 mRNAs are informative. Notice that this minimum will be unique to each observation (combination of cassette exon and cell), as it depends on the cell-specific coverage rate, and the cell and exon specific observed 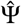.

To determine if the selected splicing observations reveal the covariance structure of the individual cell populations, we used the Chen et al. [15], Trapnell et al. [33], and Song et al. [5], datasets. We selected the exons for which we accepted observations in at least 50% of the cells. To determine the covariance structure of the data, we calculated the covariance matrix of the cells using the ormalized 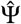 observations (due to the high variability of single cell data, we set the maximum and minimum z-scores as 3 and −3 respectively). We determined the percent of variability captured by the first *k* eigenvalues as follows:

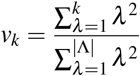

where *v*_*k*_ is the percent of variability captured by the first *k* eigenvalues, and Λ is the set of all eigenvalues of the covariance matrix. We determined the percent of variability captured by the first *k* ∈ {1, …, 10} eigenvalues in the three datasets. We compared the structure of the data with our filtering criteria versus filtering the data based on mRNA counts only, and based on reads only. For filtering with mRNAs only, we kept the splicing observations of cassette exons that fall in genes with at least 10 estimated mRNA molecules. For filtering based on reads only, we kept the observations that were covered by at least 10 splice junction reads. For each filtering approach, we selected the exons for which we accepted observations in at least 50% of the cells, and repeated our covariance structure analysis for the first *k* ∈ {1, …, 10} eigenvalues.

### Theoretical analysis of the observed 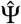 rate with limited capture

mRNA molecules are captured at a limited rate, approximated in some instances at 10% of the molecules in the cell. Under the assumption of uniform sampling of transcripts and isoforms, and assuming the only nuisance factor is the limited capture rate, we formalize the probability for observing a splicing ratio 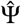. We start by specifying this probability, assuming that we know the total number of transcripts from the respective gene in the cell (*m*), the real splicing rate Ψ and the number of captured molecules *r* (assuming that for any capture molecule we know if it includes the exon or not). In that case:

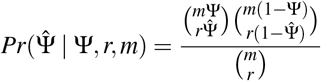

Note that for this calculation, the capture efficiency (*c*) is not needed, since we assume that we know *m* and *r*. For a more useful analysis, we will next assume that only one of these variables is not known (starting with *m* and then *r*).

In a more realistic scenario, *r* and *c* can be estimated (e.g., using Census), while *m* remains unknown. We can therefore marginalize *m* to calculate:

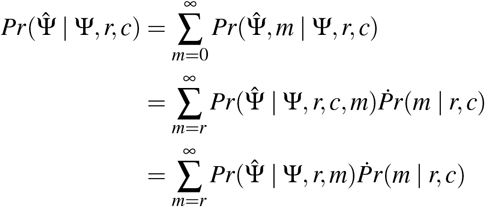

To estimate *Pr*(*m* | *r, c*) we note the following:

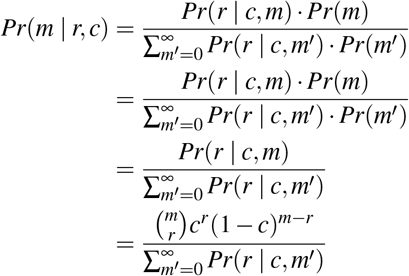

where we model the probability of capturing *r* mRNA molecules as a Binomial sample from *m* with probability *c*. Note that the third transition is done under the assumption of a uniform prior on *m*.

To compute the denominator, we expand:

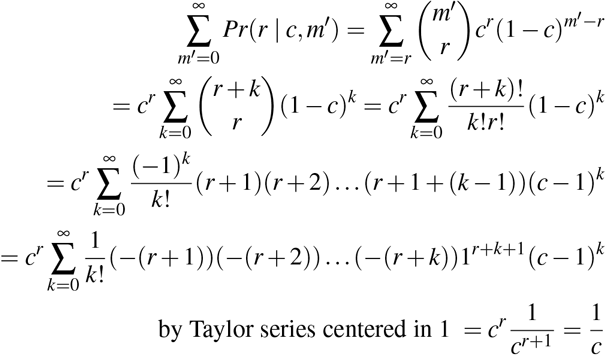

Thus

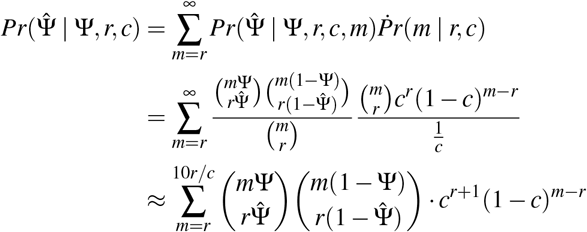

In the last equation, we estimate the sum going only up to a large value of *m*, since its posterior probability diminishes. In expectation *m ≈ r/c*. We use ten times this value as the maximum.

We can use this equation to estimate the expected proportion of binary 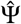 observations that is expected when we observe only *r* junctions from a splicing event with a given true rate Ψ (Supplementary Figure 2a). We can also estimate the chance to have an empirical 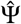 that is at least within a certain delta (in absolute terms) from the real Ψ. Namely we can estimate 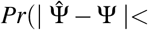 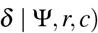 (Supplementary Figure 2b).

In another calculation of interest, one can ask how many mRNA molecules should a gene have in a cell in order to correctly estimate the splicing rate, under a limited capture efficiency *c*. To estimate it, we denote by *d* a binary variable indicating that the gene has been detected (i.e., *r* > 0) and marginalize *r* in the following way:

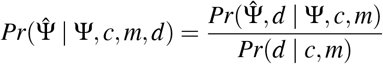

Where

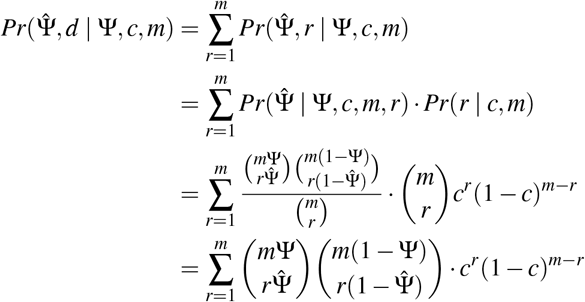

and

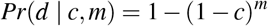

We use this in Supplementary Figure S2c to plot the chances to see only one isoform (binary 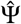) for a fixed Ψ (set to 0.5) as a function of the number of molecules present in the cell (*m*).

### Probabilistic simulator of splicing in single cell data

1) Biological process. We simulate the expression of 500 genes in 300 cells using SymSim, an *in-silico* simulator of gene expression in single cells [20]; the expression of gene *g* in cell *i* is annotated as *X*_*i*_. We simulate one cassette exon *j* for each gene *g*. For each exon *j* in cell *i*, we simulate an underlying splicing distribution of a cassette exon *j* Ψ_*i j*_ as a Beta distribution with exon-specific parameters *α* _*j*_ and *β* _*j*_. The splicing of *j* in *i* is simulated as a Binomial sampling from *X*_*ig*_ with probability Ψ_*ij*_. 2) Technical process. We simulate the capture, fragmentation and sequencing of each transcript using a modified version of SymSim’s `True2ObservedCounts` function and a random vector of transcript lengths. Finally, we subsample the obtained reads based on the transcript length in order to simulate the coverage of informative splice junctions.

We simulate the splicing of cassette exons in a set of genes *G* expressed in a population of cells *N*. For each gene *g ∈ G*, we simulate the splicing of one cassete exon *j*. The inclusion of *j* forms the isoform *j*_*A*_, while the exclusion of the exons forms the isoform *j*_*B*_. The production of mRNA molecules from *j*_*A*_ in a single cell *i ∈ N* is determined by the total expression of *g*, and by the action of the splicing machinery of *i*.

#### 1. Biological process

##### Splicing from pre-mRNA transcripts in individual cells

For each alternatively spliced gene *g* in a cell *i*, we simulate the expression of *g* and the splicing of its cassette exon.

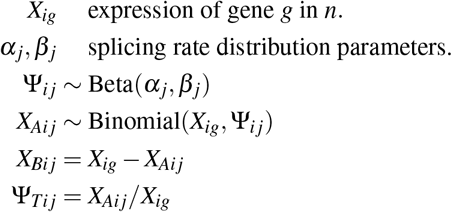

*X_ig_* represents the total number of pre-mRNA transcribed from each gene across all cells. We simulate the total counts using SymSim, an *in-silico* approach for simulation of single cell gene expression by accounting for the biological sources of variation [20]. Ψ_*ij*_ referred to as the underlying splicing rate, is the probability of splicing-in the cassette exon *j* of gene *g* in cell *i*. Notice that in this simulation, each gene *g* only has one cassette exon In a biological context, Ψ_*ij*_ would be determined by intrinsic attributes of the cassette exon inherent of *g* (e.g., sequence, secondary structure, binding sites), and by the profile of splicing factors expressed in *n*. *X*_*Aij*_ and *X*_*Bij*_ are respectively the counts of mRNA molecules from isoforms *g_A_* and *g_B_* in *i*. Notice that *X*_*Aij*_ is a random binomial sample from the total number of expressed pre-mRNA molecules of *g* in *i* with a probability Ψ_*ij*_, as it has been modeled before [10, 28, 34, 35]. Ψ_*Tij*_ is the true isoform ratio that include cassette exon *j* in cell *i*, obtained as the proportion of molecules of gene *g* that include the cassette exon *j*.

The distribution of Ψ_*ij*_ across all cells *i ∈ N* is modeled as a Beta distribution, which has been used in previous studies of single cell splicing [5, 11, 21]. In this model, the distribution is determined by the parameters *α*_*j*_, *β*_*j*_ ∈ (0, ∞). The values of these parameter determine the distribution of Ψ_*ij*_ across all cells as follows:

- Unimodal with intermediate mode if *α*_*j*_, *β*_*j*_ > 1.
- Unimodal with mode 1 if 0 < *α*_*j*_ < 1 ≤ *β*_*j*_.
- Unimodal with mode 0 if 0 < *β*_*j*_ < 1 ≤ *α*_*j*_.
- Bimodal with modes 0 and 1 if 0 < *α*_*j*_, *β*_*j*_,< 1.
- Uniform if *α*_*j*_ = *β*_*j*_ = 1.

Notice that 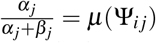. By controlling the *α*_*j*_, *β*_*j*_ parameters we can compare the biological underlying distribution of the exon splicing rate with the observed distribution of Ψ inferred from single cell RNA-seq data.

To compare the results of our simulations under the bimodal and unimodal models of splicing, we considered two competing distributions for the underlying splicing distributions of each cassette exon *j* in cell *i*:

- Unimodal splicing:

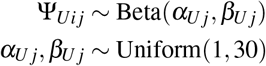
- Bimodal splicing:

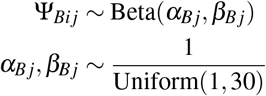

To simulate a realistic scenario, for both the unimodal and bimodal models, we simulated additional exons that are consistently included or consistently excluded. For the consistently included exons, we sampled the Beta parameters as

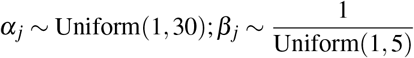

For the consistently excluded exons, we sampled the Beta pa-rameters as

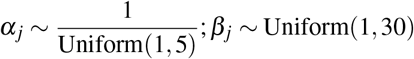

For the bimodal simulations shown in Figure 2, we simulated 500 intermediate exons with bimodal Beta distributions, 500 consistently excluded exons, and 500 consistently included exons. For the unimodal simulations, we simulated 500 intermediate exons with unimodal Beta distributions, 500 consistently excluded exons, and 500 consistently included exons.

#### 2. Technical process

##### mRNA capture into cDNA

After simulating the production of mRNAs of distinct isoforms in single cells, we simulate the process of capture and sequencing of mRNA molecules from *X*_*Aij*_ and *X_Bij_*.

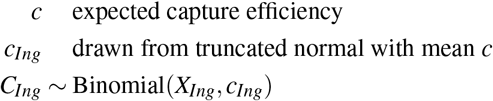

The process of mRNA capture is simulated using SymSim. *C*_*Ing*_ is the number of mRNA molecules of isoform *I* ∈ {*g*_*A*_, *g*_*B*_} for gene *g* on cell *n*. This number is sampled from the total number of molecules for isoform that are present in the cell. *c* is a parameter that determines the expected capture efficiency. *c*_*Ing*_ is the specific probability of capture of isoform *I* of gene *g* in cell *n*, which is drawn from a truncated normal distribution with mean *c*.

##### RNA sequencing

Sequencing is also simulated using SymSim’s approach with a slight modification. Artificial length amplification bias can substantially deviate the ratio between two isoforms that come from the same gene. In order to avoid that, we sampled the amplification biases from SymSim before simulating amplification on each cell, instead of doing it only once before simulating amplification for all cells, as in the original algorithm. This was shown to successfully eliminate unwanted gene length amplification bias that is not relevant for the focus of this study.

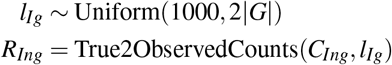

*l*_*Ig*_ is the length of isoform *I* in gene *g*, with *I* ∈ {*A, B*}. We assigned these lengths to each isoform drawing without replacement from a uniform distribution with range [1000, 2|*G*|], with |*G*| the total number of genes being simulated. *R*_*Ing*_ is the total number of observed reads from isoform *I* of gene *g* in cell *n* obtained from single cell RNA sequencing. These are obtained using the True2ObservedCounts function from SymSim with the modification previously described.

##### Splice junction coverage and observed Ψ calculation

We also simulate the down-sampling from observing only reads that overlap the splice junctions that are informative about the splicing of the cassette exon.

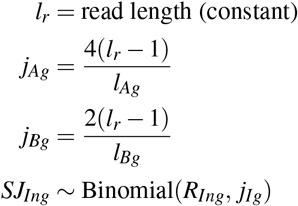

*l*_*r*_ corresponds to the read length from the sequencing process, which is assumed to be constant. *SJ*_*Ing*_ is the number of reads that cover informative splice junctions for isoform *I* ∈ {*g*_*A*_, *g*_*B*_} for gene *g* in cell *n*, which are sampled from the total number of reads covering the isoform. *j*_*Ag*_ and *j*_*Bg*_ are respectively the probabilities of a given read to cover the splice junctions informative with isoform *g*_*A*_ and *g_B_*. We assume that the distribution of reads is expected to be uniform across the transcript. Each read can be mapped to 2(*l*_*r*_ − 1) positions in the transcript that overlap one splice junction. Thus, the probability of covering one given splice junction is defined as the number of possible positions in the transcript that are informative for the splice junction, divided by the length of the transcript. *j*_*Ag*_ is the probability to map to any of the two splice junctions that are informative for isoform *g_A_*. *j*_*Bg*_ is the probability to map to one single splice junction, since there is only one junction informative for isoform *g_B_*.

Finally, the observed Ψ is calculated as:

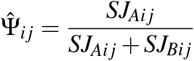

### Simulator variants for studying sources of variation

#### Gene expression and underlying Ψ

We tested the effect of the interplay between gene expression and the ratio of isoforms that contain the cassette exon on the observed distribution of Ψ. For this test, we simulated a population of 300 single cells with 500 genes, indexed 1 to 500. For every cell *i*, the expression of gene *g* is fixed as *X*_*ig*_ = *g*, where *g* ∈ {1, 2, …, 500}. This is, every gene had a different level of expression, and the expression of every individual gene was constant across all cells. For each simulation, we fixed the underlying splicing rate of all cassette exons across all cells. This is, for each cassette exon *j* of gene *g*, in every cell, we set Ψ_*ij*_ = constant. We ran the simulator with different underlying splicing rates, with Ψ_*ij*_ ∈ {0.01, 0.02, …, 0.5}. For every simulation we used an average capture efficiency *c* = 0.1. We ran 50 simulations for every fixed Ψ_*ij*_ value. For every fixed Ψ_*ij*_ and for every fixed expression level *g*, we took the average proportion of cells with binary values for the observed 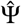. This is, we reported:

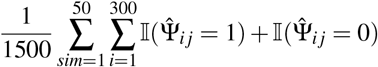

#### Gene expression and capture efficiency

We tested the effect of capture efficiency in Ψ observations. To minimize the effect of the underlying Ψ in the simulations, in this analysis we fixed the true splicing rate of all exons to Ψ_*Tng*_ = 0.5 (we achieved this by setting 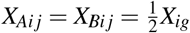). We ran simulations for each possible value for the average capture efficiency in *c* ∈ {0.01, 0.011, …, 0.1}. For each tested average capture efficiency rate, we ranked the alternative splicing events by the number of reads that cover the informative splice junctions. For each alternative event, we observed the proportion of cells that present only one type of isoform (either including the cassette exon or excluding it, but not both).

#### cDNA preparation process

The results so far suggest that the process of capturing mRNA molecules into cDNA is a limiting factor in the estimation of Ψ in single cells. Given that capture efficiency has a large effect in the distortion of Ψ distributions, we asked if other parameters that influence the preparation of cDNA libraries also affect Ψ, namely, cDNA amplification. The single cell RNA-seq datasets that we analyzed vary the number of PCR cycles after reverse transcription for cDNA library amplification. We asked if the number of PCR amplification cycles and the amplification efficiency of the experiments affect the distirbution of Ψ. To test this possibility, we ran simulations with a fixed the true splicing rate to Ψ_*Tng*_ = 0.5. We set the number of cDNA PCR cycles as *cDNA* ∈ {10, 11, …, 25}, and the PCR amplification efficiency as *η* ∈ {0.5, 0.55, …, 0.95}. We used these parameters as inputs of the modified SymSim sequencing simulation s tep. For each possible combination of the *cDNA* and *η* parameters, we ran 50 simulations and obtained the average binarity of the data. We found that incleasing the number of PCR amplification cycles also increases the proportion of cells in which we observe binary Ψ. Interestingly, PCR amplification efficiency only has a subtle effect in the porportion of cells with binary Ψ, in which higher amplification efficiency values decrease the proportion of binary observations.

#### cDNA fragmentation and fragment amplification

In contrast, changing the number of PCR cycles for fragment amplification did not have a large impact on the estimation of the original Ψ. This suggests that the bottleneck for detecting splicing isoforms in single cells comes mostly from the process of cDNA capture and gene expression, not from fragmentation and amplification (Supplementary Figure S2f,g).

## Acknowledgements

We thank Don Rio and Chenling Xu for discussions that inspired this analysis, and Nicholas Ingolia for critical comments on the manuscript. C.F.B. was supported by the UC MEXUS-CONACYT doctoral fellowship.

---

**Figure S1:**
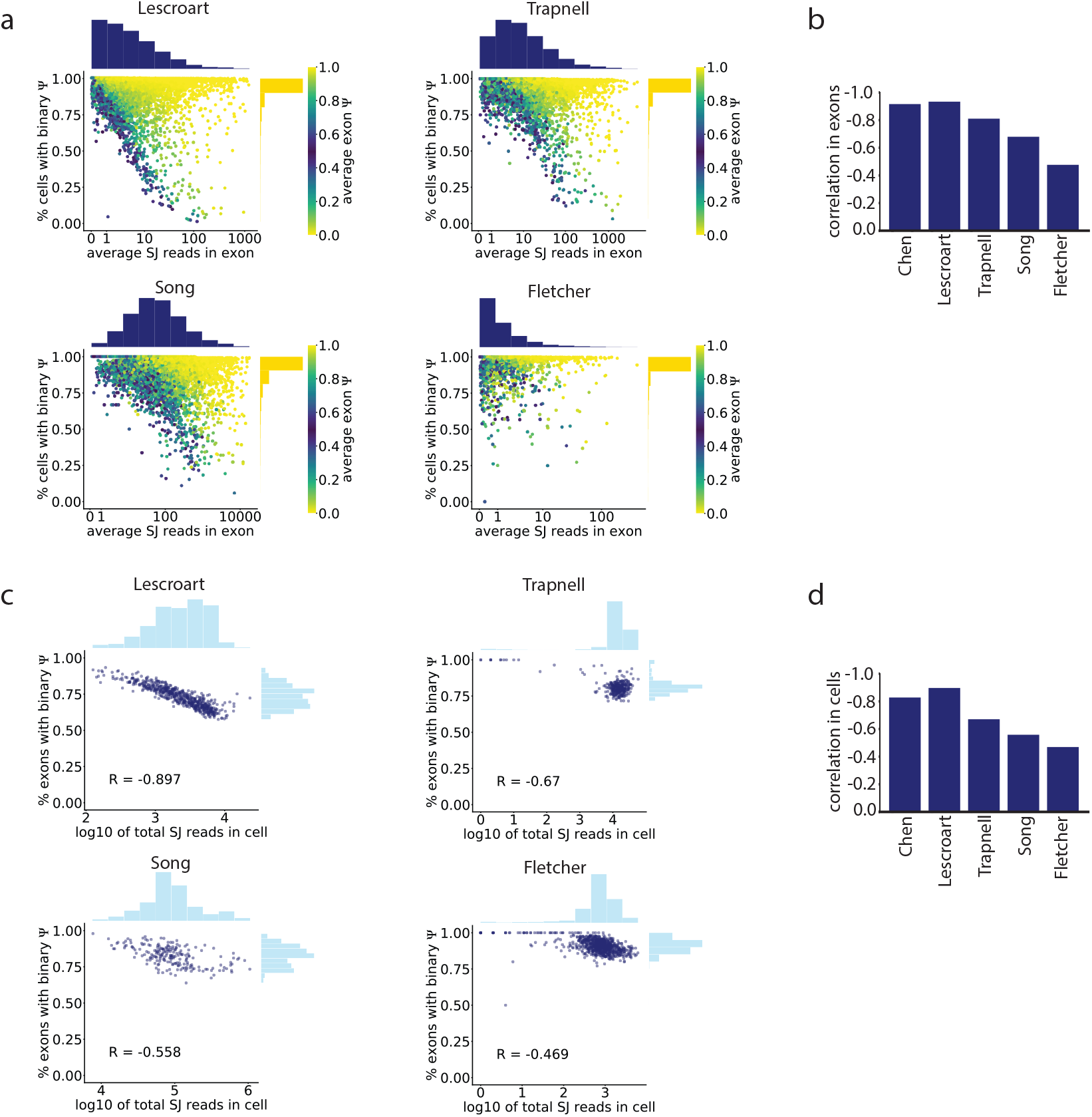
a) As in Figure 1c, the splice junction read coverage of an intermediate exon was anti-correlated with the proportion of cells in which it shows binary 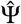 in all datasets analyzed. b) Pearson correlation score between read coverage and proportion of binary observations of each cassette exon across five scRNA-seq datasets. c) As in Figure 1d, the total number of splice junction reads in a cell was inversely proportional to the fraction of exons that have binary values in the cell in all datasets analyzed. d) Pearson correlation score between the total number of captured reads in each cell and the proportion of cassette exons with binary 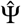 across five scRNA-seq datasets.

**Figure S2:**
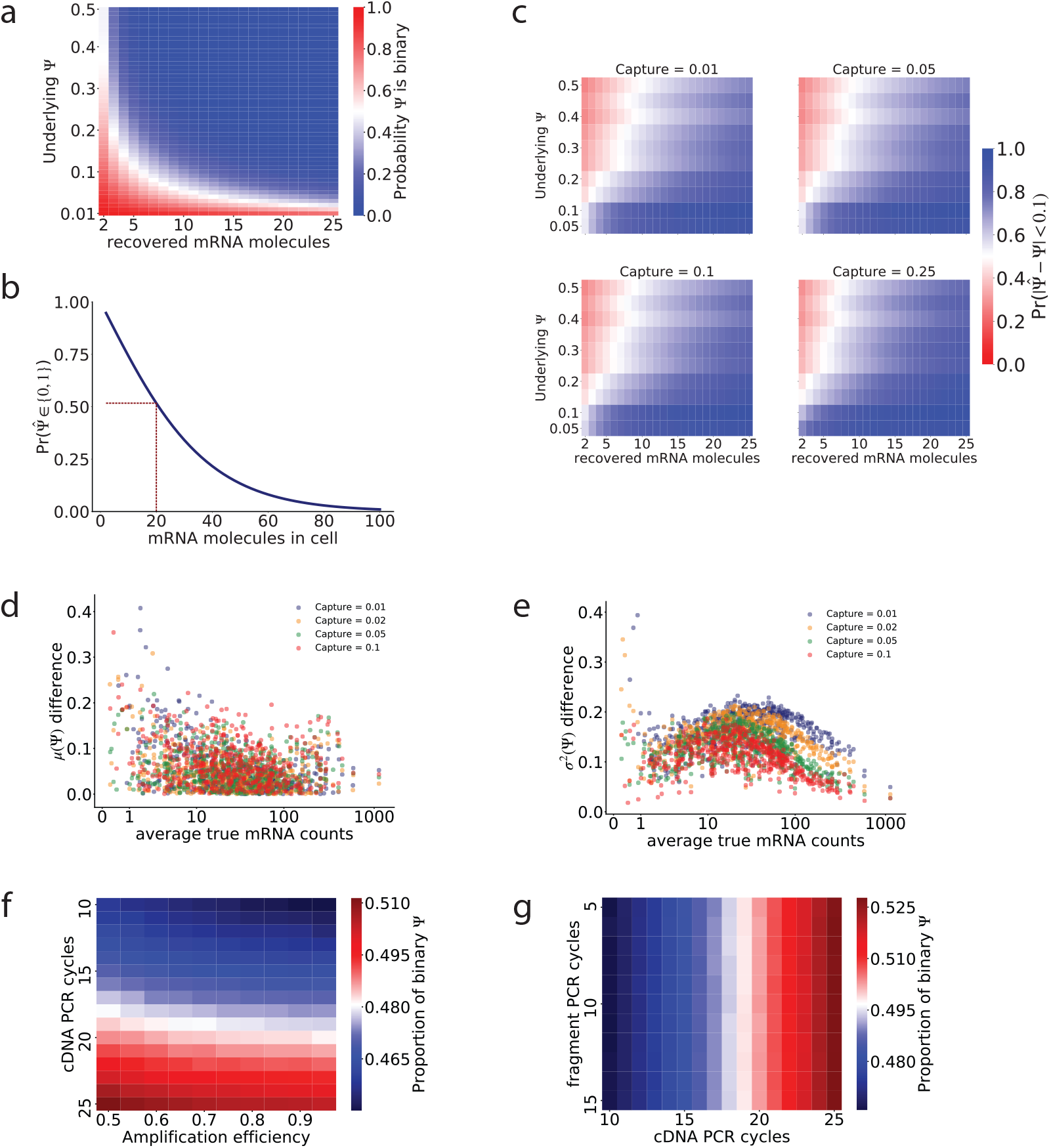
a) Theoretical likelihood of capturing only mRNAs representing one isoform of an alternatively spliced gene in a single cell, determined by the total number of mRNAs in the cell, the Ψ of the isoforms, and a 10% capture efficiency. b) Probability of observing a binary 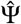 given the number of mRNA molecules of that gene present in the cell. c) The probability of having an observed 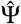 within 0.1 of the underlying Ψ. c) Difference between the average true Ψ (fraction of mRNAs selected to include the skipped exon), and the average observed 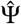 determined by splice junction reads in the intermediate exons from the simulations. The difference in average Ψ decreases as there are more initial mRNAs, but it is not affected by the capture efficiency. d) Difference between the variance in the true Ψ, and the variance in the observed 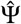 in the intermediate exons from the simulations. The difference in average Ψ decreases as there are more initial mRNAs. The difference in variance is larger as the capture efficiency is lower. e) Effect of the number of cDNA PCR cycles and amplification efficiency on the proportion of binary observations in our simulations, with the underlying Ψ fixed at 0.5 and capture efficiency at 10%. The effect of both parameters is modest compared with the effect of capture efficiency, yet clear. This result suggests that compensating the low starting material in single cells with excessive cDNA PCR amplification might worsen the distortion in splicing observations. f) Effect of the number of cDNA PCR amplification cycles and fragment amplification cycles in the proportion of binary observations.

**Figure S3:**
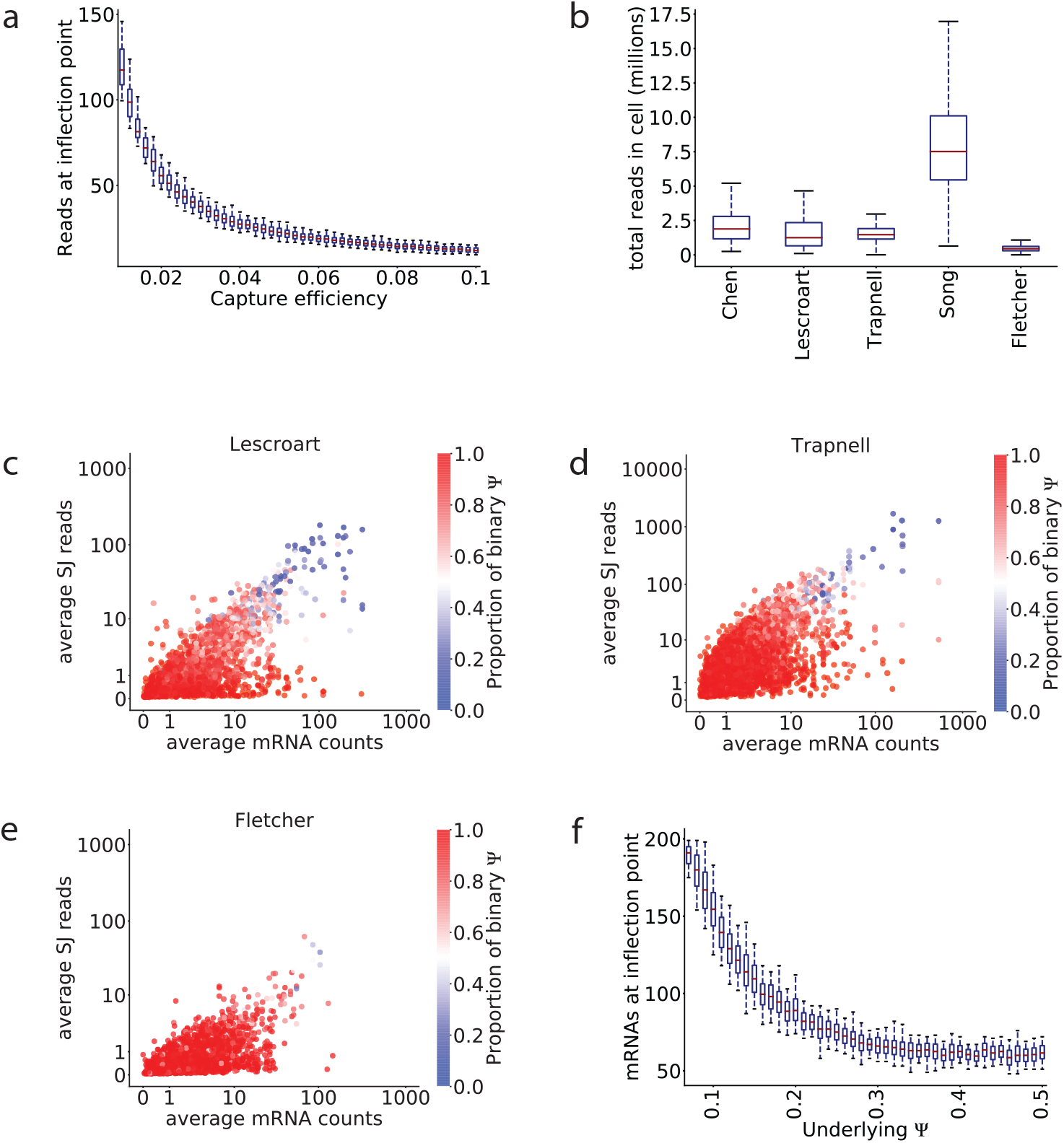
a) Average number of splice junction reads required to have a 50% likelihood of observing both isoforms in the simulations when the underlying Ψ is fixed to 0.5, under different capture efficiency values and a fixed expected sequencing library size. The average number of reads required to detect both isoforms increases as the capture efficiency decreases. These results come from the simulations shown in Figure 2j. b) Total number of mapped reads per cell for each dataset. c-e) Comparison of the average number of recovered mRNAs (from the Census estimate) and the average number of splice junction reads for each exon in the c) Lescroart et al [36] dataset, d) Trapnell et al [33] dataset, and e) Fletcher et al [37] dataset. f) Total number of mRNAs required to have a 50% likelihood to observe two isoforms in the simulations, for exons with different Ψ at a capture efficiency of 10%. Based on these results, and considering the 10% capture efficiency in the simulations, it appears that a minimum of 10 captured mRNA molecules per observation would result in a good quality splicing observation.

